# DeepRNA-Twist : Language Model guided RNA Torsion Angle Prediction with Attention-Inception Network

**DOI:** 10.1101/2024.10.24.619978

**Authors:** Abrar Rahman Abir, Md Toki Tahmid, Rafiqul Islam Rayan, M Saifur Rahman

## Abstract

RNA torsion and pseudo-torsion angles are critical in determining the three-dimensional conformation of RNA molecules, which in turn governs their biological functions. However, current methods are limited by RNA’s structural complexity and flexibility, as it can adopt multiple conformations, with experimental techniques being costly and computational approaches struggling to capture the intricate sequence dependencies needed for accurate predictions. To address these challenges, we introduce DeepRNA-Twist, a novel deep learning framework designed to predict RNA torsion and pseudo-torsion angles directly from sequence. DeepRNA-Twist utilizes RNA language model embeddings, which provides rich, context-aware feature representations of RNA sequences. Additionally, it introduces 2A3IDC module (**A**ttention **A**ugmented **I**nception **I**nside **I**nception with **D**ilated **C**NN), combining inception networks with dilated convolutions and multi-head attention mechanism. The dilated convolutions capture long-range dependencies in the sequence without requiring a large number of parameters, while the multi-head attention mechanism enhances the model’s ability to focus on both local and global structural features simultaneously. DeepRNA-Twist was rigorously evaluated on benchmark datasets, including RNA-Puzzles, CASP-RNA, and SPOT-RNA-1D, and demonstrated significant improvements over existing methods, achieving state-of-the-art accuracy. Source code is available at https://github.com/abrarrahmanabir/DeepRNA-Twist

## Introduction

RNA molecules are fundamental to numerous biological processes and their functionality is closely linked to their three-dimensional structures, akin to proteins. The functionality of RNA critically depends on its structural configuration, highlighting the need for precise characterization of its complex three-dimensional conformation. Traditional experimental methods for determining RNA structure, such as NMR [1], X-ray crystallography [2], and cryo-EM [3], while reliable, are often limited by high costs and time-consuming processes.

Central to RNA’s structural complexity are its torsion angles, comprising seven specific angles (*α, β, γ, δ, ϵ, ζ* and *χ*) along the ribose-phosphate backbone. Additionally, two pseudo-torsion angles (*η* and *θ*) provide a simplified representation of the RNA backbone (Figure 1). These angles are crucial as they dictate RNA’s folding patterns and ultimately its three-dimensional shape, enabling RNA to perform diverse biological roles—from catalysis as ribozymes to regulating gene expression through various non-coding RNA mechanisms. Understanding these torsion angles enhances the capability to design therapeutic agents [4], potentially altering their function to treat diseases at a molecular level.

**Fig 1.**
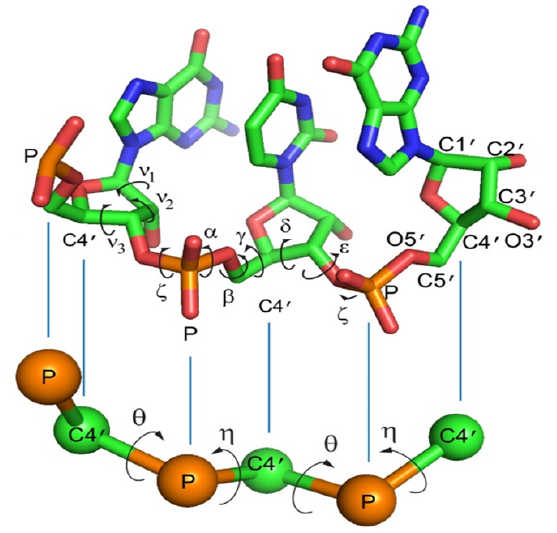
Top: A detailed view of a short RNA sequence (GUG) illustrating all atoms excluding hydrogens along with key atoms and native torsion angles labeled. **Bottom**: pseudo-torsion angles *η* and *θ* for the same sequence. From [10], Copyright 2016 The Authors under the Creative Commons Attribution 4.0 International License (https://creativecommons.org/licenses/by/4.0/).

Predicting RNA torsion angles and structures presents substantial challenges due to RNA’s inherent properties and the limitations of current methodologies. Unlike proteins, RNA’s structure is determined by a more complex system of torsion angles that contribute to its flexibility and dynamic nature, allowing multiple conformations [5]. This complexity is compounded by RNA’s engagement in non-canonical interactions such as Hoogsteen base pairing, base triples, and various loop interactions, vital for its biological functions but complicating structural predictions.

Recent advancements in protein torsion angle prediction have effectively utilized deep learning methods to enhance prediction accuracy [6, 7]. Inspired by the success of deep learning for predicting torsion angles of protein, researchers have employed various deep learning approaches to predict RNA torsion angles as well, although this area remains very underexplored. SPOT-RNA-1D [8], the first RNA backbone torsion angle prediction method, utilizes a dilated convolutional neural network to predict both torsion and pseudotorsion angles of RNA from single sequence inputs. Currently, the development of language models that can predict RNA structural features solely from sequence data is very limited. RNA-TorsionBERT [9] model leverages language model approach to predict torsional and pseudo-torsional angles directly from sequence data. Additionally, this model introduces RNA Torsion-A, a scoring function that assesses the quality of predicted RNA structures using the torsion angles generated by the model, further refining the evaluation of RNA structural predictions.

In this study, we present DeepRNA-Twist, a novel deep learning approach for predicting RNA torsion and pseudotorsion angles directly from sequence data. A key novelty of DeepRNA-Twist is that it utilizes RNA language model embeddings (RiNALMo [11]) as input features which contribute to the model’s improved performance. This novel approach to feature representation has not been explored before for RNA torsion angle prediction.

Additionally, the introduction of the 2A3IDC (**A**ttention **A**ugmented **I**nception **I**nside **I**nception with **D**ilated **C**NN) module is our another major contribution. This module combines inception network and dilated convolutional neural network with multi-head attention mechanism. The use of dilated CNN allows the model to capture a broader context and long-range dependencies in the sequence data while enhancing computational efficiency by reducing the number of parameters required to achieve a large receptive field. This enables the model to process sequences more efficiently without sacrificing performance. On the other hand, the multi-head attention mechanism further improves the model’s ability to focus on different parts of the sequence simultaneously, enhancing the capture of both local and global structural features. DeepRNA-Twist effectively captures both short and long-range interactions among nucleotides. We rigorously evaluated DeepRNA-Twist against existing leading approaches on widely recognized benchmark datasets such as RNA-Puzzles [12] and CASP-RNA [13] and SPOT-RNA-1D [8] Dataset. Our results demonstrate that DeepRNA-Twist significantly outperforms existing methods, achieving state-of-the-art accuracy in RNA torsion angle prediction.

## Materials and Methods

### Dataset

We used SPOT-RNA-1D [8] dataset for training and validation. We evaluated our DeepRNA-Twist on two independent test sets - SPOT-RNA-1D Test dataset [8] and RNA-TorsionBERT Test dataset.

#### SPOT-RNA-1D Dataset

For SPOT-RNA-1D [8], RNA structures were sourced from the Protein Data Bank (PDB), selecting those with X-ray resolutions finer than 3.5 Å as of October 3, 2020. As described by the authors of SPOT-RNA-1D [8], these structures were segmented into individual chains using Biopython, clustered with CD-HIT-EST [14] at an 80% identity threshold to form the training set, with unclustered sequences making up a noncluster set. To refine the data, BLAST-N [15] filtered out sequences with internal or cross-set similarities using an e-value cutoff of 10. The noncluster sequences were subsequently divided into a validation set (VL) and two test sets (TS1 and TS2), ensuring minimal redundancy by constructing covariance models using the INFERNAL tool’s cmbuild and cmsearch programs, applying strict e-value cutoffs to eliminate remote homologues. An additional test set (TS3) was later formed from NMR structures, processed similarly to ensure nonredundancy. The final datasets included 286 RNA chains for training, with 30 for validation, and 63, 30, and 54 for TS1, TS2, and TS3 respectively where the maximum sequence length is 418. Native torsion angles were extracted using the DSSR program [16].

#### RNA-TorsionBERT Test Dataset

RNA-TorsionBERT [9] employed a combined test set consisting of two prominent datasets: RNA Puzzles [12] and CASP-RNA [13]. This merged dataset consists of 52 structures— 40 from RNA Puzzles and 12 from CASP-RNA. The length of sequences of this dataset ranges from 27 to 512 nucleotides. Figure 4 demonstrates the distribution of all torsion and pseudo-torsion angles of both datasets.

### Feature Representation

DeepRNA-Twist takes an RNA sequence feature vector *X* = {*x*_1_, *x*_2_, …, *x*_*i*_, *x*_*i*+1_, …, *x*_*N*_ } as input, where *x*_*i*_ is the vector corresponding to the *i*-th nucleotide of that RNA. Inspired by the success of protein language models in capturing biological patterns and in downstream tasks, we used RiNALMo, an RNA language model developed by [11], to generate input features. However, DeepRNA-Twist is agnostic about the language model. RiNALMo generates a sequence of embedding vectors *X* = {*x*_1_, *x*_2_, …, *x*_*N*_ }, where each and *d*_RiNALMo_ = 1280, represent the features for *i*-th nucleotide of RNA sequence.

### DeepRNA-Twist Architecture

DeepRNA-Twist architecture is composed of several integrated modules designed to capture complex patterns and dependencies within RNA sequences. The architecture includes: Transformer Encoder Layer, Multi-head Attention module and 2A3IDC module.

#### Transformer Encoder Layer

DeepRNA-Twist incorporates a Transformer Encoder Layer [17] to process the RNA sequence feature vector *X* = {*x*_1_, *x*_2_, …, *x*_*N*_ }, where each *x*_*i*_ is a 1280-dimensional vector corresponding to the *i*-th nucleotide, provided by the RiNALMo [11]. Each Transformer Encoder Layer consists of two main components: a multi-head self-attention mechanism and a position-wise feed-forward network. The multi-head self-attention mechanism refines the representation of each nucleotide by attending to all other nucleotides in the sequence. For each nucleotide *x*_*i*_, the self-attention mechanism produces an intermediate representation **a**_*i*_ that captures contextual dependencies. Following the self-attention mechanism, each intermediate representation **a**_*i*_ is independently passed through a position-wise feed-forward network, resulting in a transformed representation **z**_*i*_. Formally, this can be expressed as:

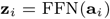

where FFN represents the feed-forward network. Each sub-layer in the encoder, including the self-attention and feed-forward networks, incorporates residual connections followed by layer normalization. This configuration stabilizes the learning process and enhances the model’s ability to capture complex dependencies across the RNA sequence. The overall transformation of *x*_*i*_ through the encoder can be summarized as:

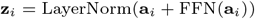

where **a**_*i*_ is obtained from the multi-head self-attention mechanism applied to *x*_*i*_. This process results in the refined representation of the intial embeddings which captures both local and global structural information of the RNA sequence.

#### Multi-head Attention Module

The multi-head attention module, denoted by Multi-HeadAttention(.), is designed to dynamically weigh the importance of different elements in the input data, adjusting the focus based on the input’s context. This module integrates a positional encoding sub-module, which provides crucial positional information to enhance the model’s ability to capture sequential relationships.

##### Positional Encoding

To encode positional information [17], the positional encoding function PE_*p*_ for a position *p* is defined as:

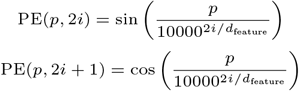

where *i* is the dimension. This allows the model to learn to attend to relative positions. The input **X** is augmented with the positional encoding, resulting in a new representation:

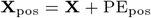

This new representation **X**_pos_ retains both the information from previous layers and the positional information of each element.

#### Projection to Query, Key, and Value

The input sequence feature vector **X** = {*x*_1_, *x*_2_, …, *x*_*N*_ } is first augmented with positional encoding to obtain **X**_pos_. This augmented input is then linearly transformed to create three matrices: Query (**Q**), Key (**K**), and Value (**V**):

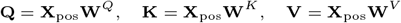

where **W**^*Q*^, **W**^*K*^, and **W**^*V*^ are learnable parameter matrices.

#### Attention Calculation

The multi-head self-attention mechanism allows the model to jointly attend to information from different representation subspaces. The input is split into *h* heads, and the attention scores are computed for each head using the scaled dot-product of **Q** and **K**, followed by scaling and masking. For the *i*-th head:

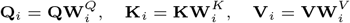

The attention scores (**A**_*i*_) are computed as:

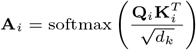

where *d*_*k*_ is the dimension of the keys. The output for each head (**O**_*i*_) is then computed by:

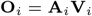

The outputs from all heads are concatenated to form the final output of the multi-head attention mechanism:

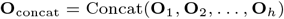

The concatenated output is then linearly transformed:

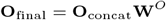

where **W**^*O*^ is a learnable parameter matrix. The resulting output is then subjected to dropout and batch normalization to enhance training stability and performance.

By integrating positional encoding and using multiple attention heads, the attention module can effectively capture both the content and positional relationships within the input, making it suitable for processing RNA sequences.

#### 2A3IDC Module

The 2A3IDC module integrates inception blocks, dilated convolutional layers, and attention mechanisms to process input data through two parallel paths, enhancing the model’s ability to capture both local and global patterns. This design builds upon the 2A3I module [18], which augments the inception-inside-inception (3I) module [19] with attention mechanisms to capture both short- and long-range interactions effectively. The 2A3I module [18] was inspired by the assembly of inception modules used in the MUFOLD-SS method for protein secondary structure prediction [19].

Each path in the 2A3IDC module begins with an inception block (Figure 2) to capture multi-scale features. The inception block consists of four parallel convolutional pathways. Given an input feature map *X*:

**Fig 2.**
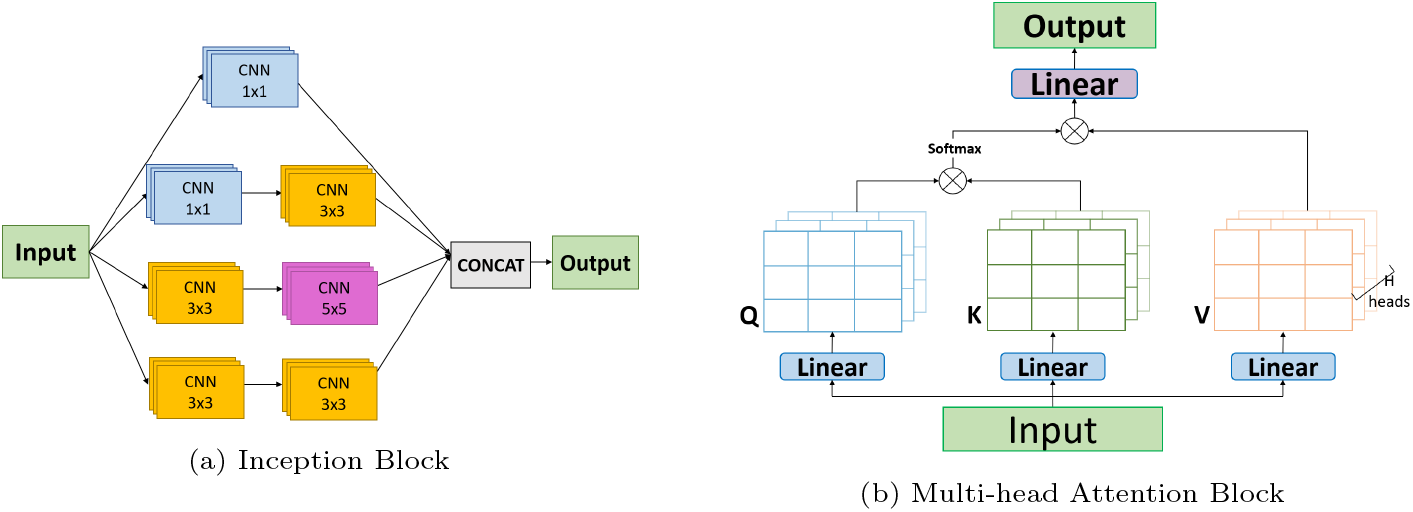
(a) Inception Block and (b) Multi-head Attention Block.

- **Path 1:** A convolutional layer with a kernel size of 1:

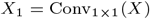
- **Path 2:** A sequence of two convolutional layers with kernel sizes of 1 and 3:

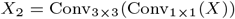
- **Path 3:** A sequence of two convolutional layers with kernel sizes of 3 and 5:

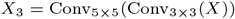
- **Path 4:** A sequence of two convolutional layers with kernel sizes of 3 and 3:

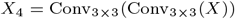

These pathways capture features at different scales, which are concatenated to form a multi-scale representation:

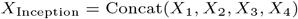

Following the inception block, each path includes a dilated convolutional layer. The dilated CNN is defined as:

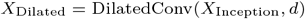

where *d* is the dilation rate, allowing the model to capture long-range dependencies efficiently without increasing the number of parameters.

The output from the dilated convolutional block is then passed through a multi-head attention module:

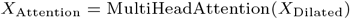

This mechanism focuses on different parts of the sequence simultaneously, enhancing the model’s ability to capture both local and global structural features.

In the 2A3IDC module, the input is processed through two parallel paths where each path consists of an inception block, followed by a dilated convolutional block, and then a multi-head attention module. We set the dilation rate as 2 and 5 for the dilated convolutional layer of the two paths respectively. Finally, the outputs from both parallel paths are concatenated and batch-normalized to form the final represen-tation:

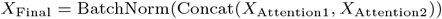

where *X*_Attention1_ and *X*_Attention2_ are the output of the first and second paths of this module.

The 2A3IDC module effectively captures and integrates both fine-grained and high-level features from the input vector, thereby enhancing the model’s prediction accuracy through a comprehensive analysis of contextual and spatial information.

#### Overall Pipeline

DeepRNA-Twist architecture predicts RNA torsion angles by integrating different modules mentioned above to capture complex patterns in the input sequence. The full pipeline of DeepRNA-Twist is shown in Figure 3. It starts with a transformer encoder that processes input RNA feature vectors generated by RiNALMo [11], followed by two consecutive 2A3IDC blocks that combine inception blocks and attention modules to capture both local and long-range dependencies. The output is passed through a 1D convolutional layer. Finally, after passing the output from an attention module, a dense layer with 18 regression nodes and tanh activation predicts the sine and cosine values of the 9 RNA torsion angles. The loss function used to train the model is the Mean Squared Error (MSE) between the predicted and true sine and cosine values of the torsion angles across all nucleotides and RNAs.

**Fig 3.**
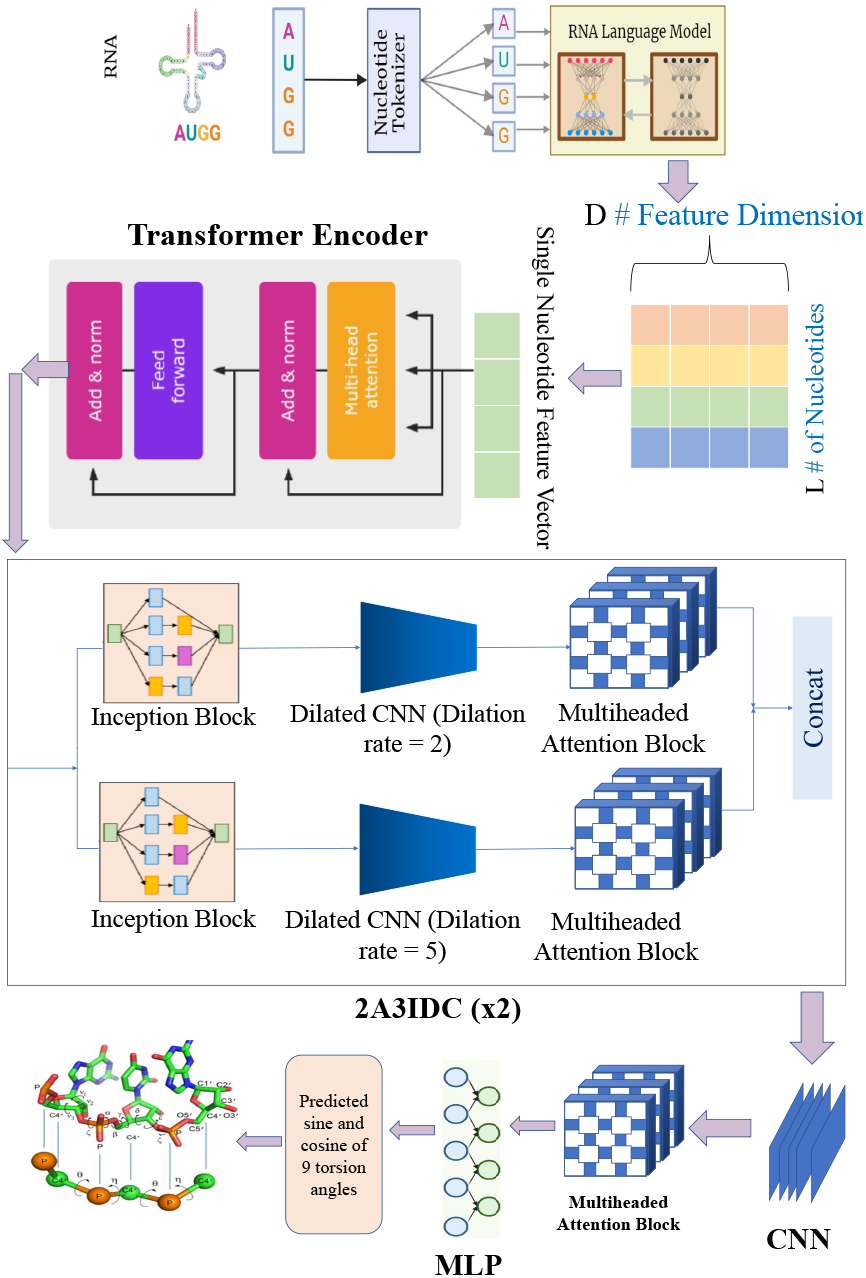
DeepRNA-Twist Architecture. The diagram of the RNA 3D structure is from [10], Copyright 2016 The Authors under the Creative Commons Attribution 4.0 International License (https://creativecommons.org/licenses/by/4.0/).

**Fig 4.**
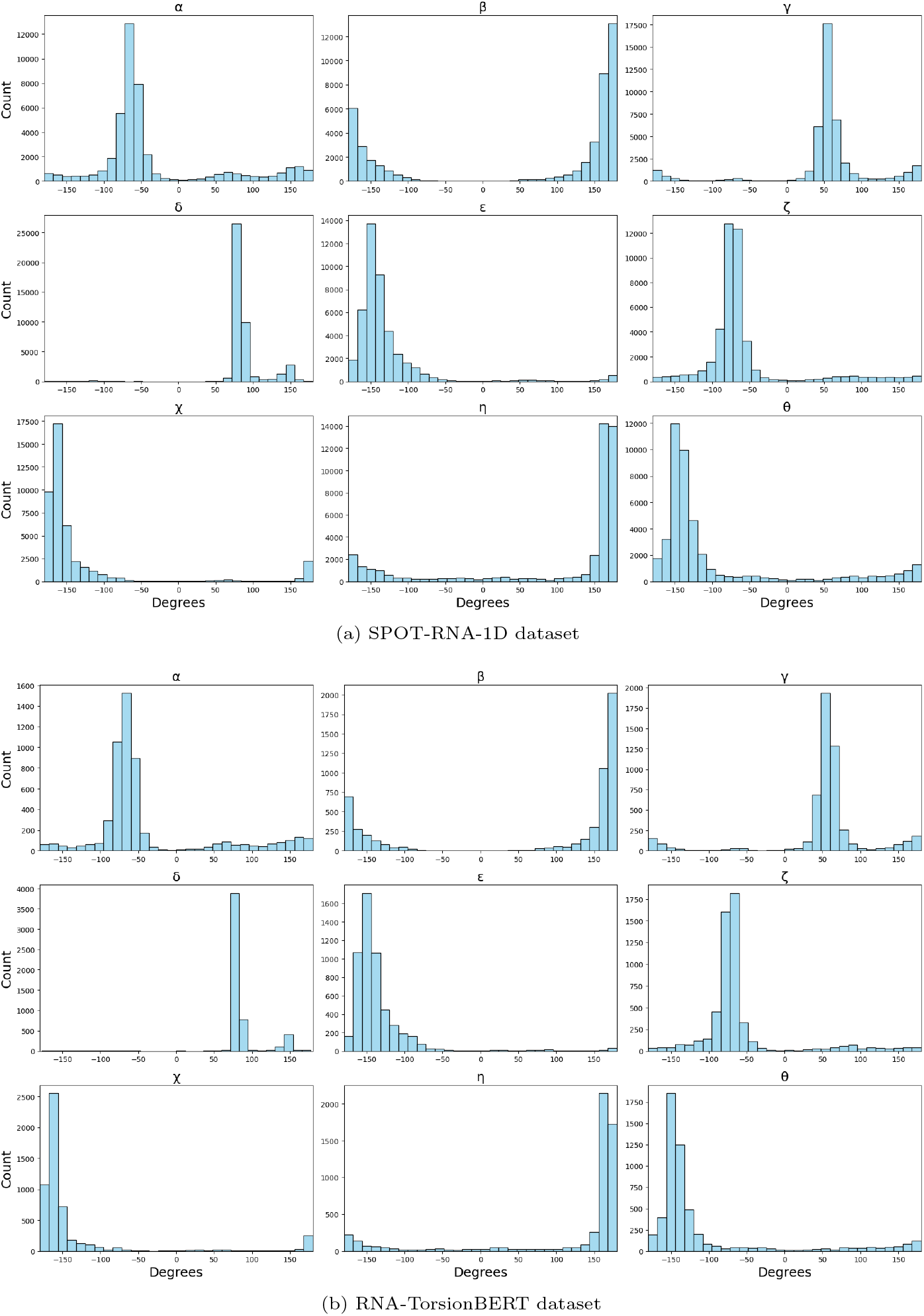
Distribution of all torsion and pseudo-torsion angles for (a) SPOT-RNA-1D dataset and (b) RNA-TorsionBERT dataset.

Let *N* be the number of RNAs, *L* be the number of nucleotides per RNA, and *A* = 9 be the number of torsion angles per nucleotide. Each angle has both sine and cosine values, leading to 2*A* = 18 values per nucleotide. The MSE loss function is defined as:

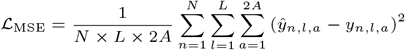

where *y*_*n,l,a*_ and ŷ_*n,l,a*_ are the true and predicted sine or cosine values corresponding to the torsion angles for the *l*-th nucleotide in the *n*-th RNA. Here, *a* ranges from 1 to 18, indexing the sine and cosine values for each of the 9 torsion angles. We trained DeepRNA-Twist for 120 epochs using the Adam optimizer with a learning rate of 0.0001.

## Results

We evaluated our model on test datasets of SPOT-RNA-1D [8] and RNA-TorsionBERT [9].

### Evaluation Metric

The predictive performance of RNA torsion angle is quantitatively assessed using the Mean Absolute Error (MAE). MAE is calculated for each torsion angle across all nucleotides in the sequence, considering the periodicity of the angles. The MAE for a specific torsion angle *θ* is defined as follows:

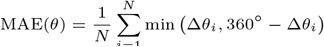

where Δ*θ*_*i*_ = |*θ*_pred,*i*_ − *θ*_true,*i*_| is the absolute difference between the predicted angle *θ*_pred,*i*_ and the true experimentally determined angle *θ*_true,*i*_ and *N* is the total number of nucleotides in the RNA sequence for which torsion angles are predicted.

### Performance on SPOT-RNA-1D dataset

DeepRNA-Twist was trained on train dataset of SPOT-RNA-1D [8] and validated using validation dataset VL. Table 2 shows the performance comparison between DeepRNA-Twist and SPOT-RNA-1D on VL and three independent test sets TS1, TS2 and TS3. In the validation set (VL), ourc model demonstrated a noticeable improvement over SPOT-RNA-1D, showing percentage reductions in MAE that ranged from approximately 10% to 15% across most torsion angles. Particularly, the reductions for the angles *α, η*, and *γ* were among the most significant. This trend of enhancement is almost consistent across the test sets, indicating robustness in handling both standard and pseudo-torsion angles. TS1 displayed similarly substantial improvements with reductions particularly pronounced for the *α* and *θ* angles. In TS2, our model excelled particularly in the prediction of *γ* and *θ*, achieving more than a 15% improvement in MAE compared to SPOT-RNA-1D. TS3, derived from structurally diverse NMR datasets, showed our model’s adaptability with significant reductions in MAE for *β* and *γ*. The performance in this set validates our model’s utility in handling RNA datasets generated from different experimental techniques.

**Table 1.** Performance comparison of DeepRNA-Twist and SPOT-RNA-1D according to MAE (Mean Absolute Error) in degrees based on various pairing interactions on SPOT-RNA-1D test sets.

| (a) TS1 |  |  |  |  |  |  |
| --- | --- | --- | --- | --- | --- | --- |
| Angles | Unpaired<br>20.92% | Lone pairs<br>15.85% | Pseudoknots<br>5.80% | Multiplets<br>7.42% | Noncanonical pairs<br>18.11% | Canonical nested pairs<br>49.84% |
| $\alpha$ | 59.55 (62.05) | 51.26 (54.26) | 51.16 (54.16) | 46.92 (49.92) | 44.24 (47.24) | 32.02 (35.02) |
| $\beta$ | 25.93 (28.43) | 24.96 (27.96) | 25.32 (28.32) | 24.52 (27.52) | 21.14 (24.14) | 15.32 (18.32) |
| $\gamma$ | 36.26 (39.76) | 37.11 (40.61) | 35.42 (38.42) | 37.73 (40.73) | 32.08 (35.08) | 26.17 (29.17) |
| $\delta$ | 17.53 (20.53) | 17.93 (20.93) | 19.03 (22.03) | 14.97 (16.97) | 14.45 (16.45) | 8.59 (10.59) |
| $\epsilon$ | 25.67 (29.17) | 25.82 (28.32) | 22.76 (25.76) | 22.52 (24.52) | 20.42 (22.42) | 13.46 (15.46) |
| $\zeta$ | 49.10 (54.10) | 47.14 (51.64) | 43.95 (47.95) | 33.84 (36.84) | 35.44 (38.44) | 17.63 (19.63) |
| $\chi$ | 28.33 (30.83) | 24.18 (27.18) | 18.91 (21.91) | 19.19 (22.19) | 20.77 (22.77) | 11.43 (13.43) |
| $\eta$ | 51.08 (54.58) | 39.26 (42.76) | 37.54 (40.54) | 32.84 (34.84) | 32.87 (34.87) | 16.24 (18.24) |
| $\theta$ | 51.20 (56.70) | 40.52 (44.52) | 39.15 (42.15) | 31.80 (33.80) | 36.83 (39.83) | 18.02 (20.02) |
| (b) TS2 |  |  |  |  |  |  |
| Angles | Unpaired<br>20.54% | Lone pairs<br>15.76% | Pseudoknots<br>6.63% | Multiplets<br>6.93% | Noncanonical pairs<br>15.89% | Canonical nested pairs<br>51.68% |
| $\alpha$ | 57.26 (60.26) | 50.62 (53.62) | 54.40 (57.40) | 47.43 (50.43) | 41.32 (44.32) | 25.03 (28.03) |
| $\beta$ | 21.49 (24.49) | 23.17 (26.17) | 26.72 (28.72) | 20.69 (22.69) | 17.78 (19.78) | 13.14 (15.14) |
| $\gamma$ | 35.89 (37.89) | 36.49 (39.49) | 32.46 (35.46) | 34.66 (36.66) | 29.99 (31.99) | 21.91 (23.91) |
| $\delta$ | 19.74 (22.74) | 20.89 (23.89) | 20.77 (23.77) | 18.13 (21.13) | 17.08 (20.08) | 9.43 (11.43) |
| $\epsilon$ | 24.92 (26.92) | 21.20 (24.20) | 16.43 (18.43) | 16.20 (18.20) | 17.43 (19.43) | 11.30 (12.30) |
| $\zeta$ | 45.34 (48.34) | 47.20 (50.20) | 30.74 (32.74) | 38.04 (41.04) | 40.14 (43.14) | 13.73 (14.73) |
| $\chi$ | 25.67 (27.67) | 25.00 (28.00) | 18.06 (21.06) | 21.32 (24.32) | 21.74 (23.74) | 10.77 (12.77) |
| $\eta$ | 49.92 (52.92) | 36.04 (39.04) | 39.38 (42.38) | 31.45 (33.45) | 30.03 (32.03) | 14.01 (16.01) |
| $\theta$ | 49.40 (52.40) | 41.06 (44.06) | 32.54 (35.54) | 31.03 (34.03) | 34.05 (37.05) | 15.21 (16.21) |
| (c) TS3 |  |  |  |  |  |  |
| Angles | Unpaired<br>18.12% | Lone pairs<br>9.20% | Pseudoknots<br>3.27% | Multiplets<br>6.41% | Noncanonical pairs<br>13.46% | Canonical nested pairs<br>60.14% |
| $\alpha$ | 64.82 (68.82) | 48.32 (51.32) | 49.06 (52.06) | 43.16 (46.16) | 49.68 (52.68) | 22.44 (25.44) |
| $\beta$ | 31.74 (33.74) | 29.53 (31.53) | 29.42 (31.42) | 27.95 (29.95) | 26.72 (28.72) | 13.40 (15.40) |
| $\gamma$ | 54.96 (58.96) | 47.84 (50.84) | 30.42 (32.42) | 43.50 (46.50) | 41.05 (44.05) | 22.61 (24.61) |
| $\delta$ | 24.39 (27.39) | 15.84 (17.84) | 16.67 (18.67) | 13.22 (16.22) | 13.20 (15.20) | 8.75 (10.75) |
| $\epsilon$ | 30.83 (32.83) | 25.01 (27.01) | 19.40 (21.40) | 22.73 (24.73) | 21.94 (22.94) | 17.68 (18.68) |
| $\zeta$ | 58.55 (61.55) | 37.07 (40.07) | 27.20 (29.20) | 33.15 (35.15) | 33.95 (34.95) | 14.99 (15.99) |
| $\chi$ | 25.66 (27.66) | 21.59 (23.59) | 13.42 (15.42) | 18.40 (20.40) | 19.01 (21.01) | 11.77 (12.77) |
| $\eta$ | 51.43 (54.43) | 28.11 (30.11) | 30.02 (32.02) | 26.60 (28.60) | 25.14 (27.14) | 14.01 (16.01) |
| $\theta$ | 55.84 (58.84) | 34.86 (36.86) | 29.55 (31.55) | 27.70 (29.70) | 25.10 (27.10) | 15.21 (16.21) |

Notes: Values within parentheses are the mean absolute error (MAE) obtained by the SPOT-RNA-1D model and reported from [8] and those without parentheses are from DeepRNA-Twist.

**Table 2.** Performance comparison of DeepRNA-Twist and SPOT-RNA-1D according to MAE (Mean Absolute Error) in degrees on SPOT-RNA-1D test sets.

| Dataset | $\alpha$ | $\beta$ | $\gamma$ | $\delta$ | $\epsilon$ |
| --- | --- | --- | --- | --- | --- |
| VL | 40.32 (45.18) | 17.40 (20.58) | 29.76 (33.88) | 14.83 (17.99) | 17.12 (20.72) |
| TS1 | 39.16 (43.94) | 17.67 (21.94) | 28.77 (32.98) | 12.79 (14.61) | 16.98 (20.69) |
| TS2 | 35.71 (39.50) | 16.12 (18.92) | 24.29 (29.47) | 13.90 (16.01) | 14.44 (17.46) |
| TS3 | 34.13 (37.89) | 16.89 (21.04) | 29.54 (34.68) | 11.30 (13.83) | 19.32 (22.32) |

| Dataset | $\zeta$ | $\chi$ | $\eta$ | $\theta$ |
| --- | --- | --- | --- | --- |
| VL | 34.68 (37.50) | 20.20 (23.01) | 29.47 (33.55) | 35.93 (37.02) |
| TS1 | 29.50 (33.27) | 16.33 (19.59) | 27.12 (30.25) | 28.52 (32.91) |
| TS2 | 25.00 (28.91) | 15.56 (18.20) | 25.19 (28.14) | 24.96 (30.25) |
| TS3 | 24.17 (27.87) | 13.92 (17.01) | 22.20 (25.31) | 25.84 (27.22) |

Notes: Values within parentheses are the mean absolute error (MAE) obtained by the SPOT-RNA-1D model and reported from [8] and those without parentheses are from DeepRNA-Twist .

**Table 3.**
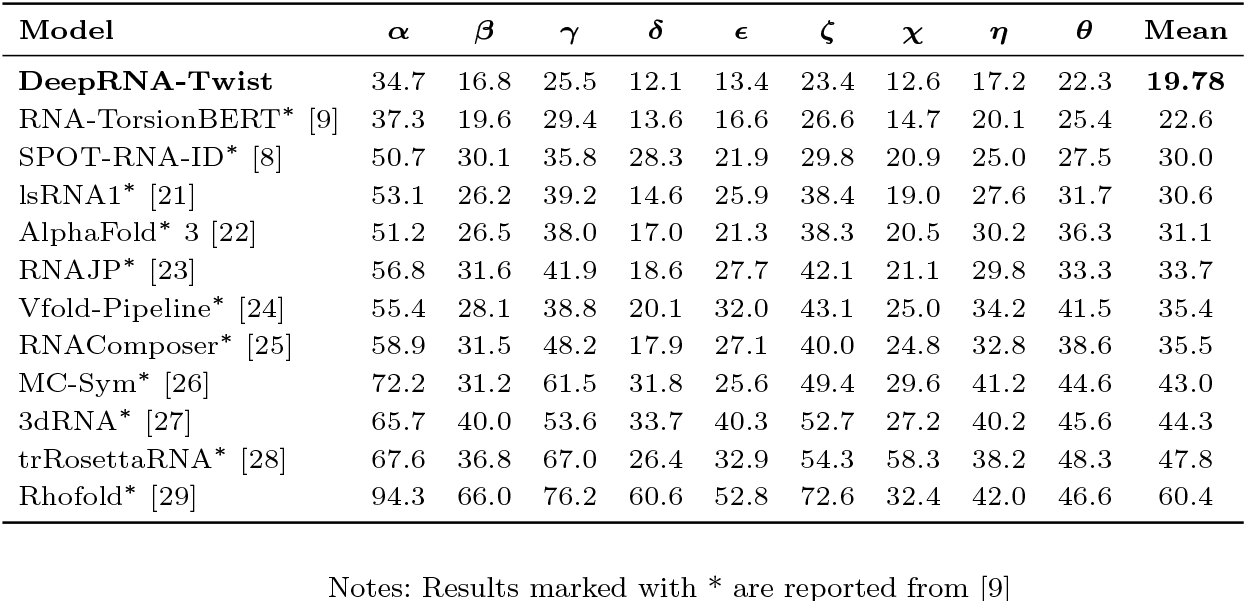
Performance comparison according to MAE (Mean Absolute Error) in degrees on RNA-TorsionBERT test sets.

The improvements in MAE were statistically significant, as evidenced by p-values from paired t-tests (Table 4). Moreover, the variation of length of RNA sequences does not lead to performance degradation as shown in Figure 5. Our analysis confirms that angles associated with narrower distributions, such as *δ, ϵ* and *χ*, are more straightforward to predict. This observation is consistent with the trends noted in SPOT-RNA-1D [8], suggesting that the distribution breadth of an angle correlates with its prediction difficulty. However, our model shows superior handling of angles that are difficult to predict due to their wider distributions, like *α, ζ*, and *θ*, by effectively reducing prediction errors. See Figure 4 for the distribution of torsion angles.

**Table 4.** P-values from one-tailed paired t-tests comparing RNA torsion angle predictions by DeepRNA-TWIST with SPOT-RNA-1D and RNA-TorsionBERT.

| Model | $\alpha$ | $\beta$ | $\gamma$ | $\delta$ | $\epsilon$ |
| --- | --- | --- | --- | --- | --- |
| SPOT-RNA-1D | 0.00038 | 0.00108 | 0.00026 | 0.00185 | 0.00019 |
| RNA-TorsionBERT | 0.0000871 | 0.0000005311 | 0.0000974 | 0.00536 | 0.00000585 |

| Model | $\zeta$ | $\chi$ | $\eta$ | $\theta$ |
| --- | --- | --- | --- | --- |
| SPOT-RNA-1D | 0.00037 | 0.00011 | 0.00050 | 0.03203 |
| RNA-TorsionBERT | 0.0000405 | 0.0004884 | 0.0000708 | 0.00000136 |

**Fig 5.**
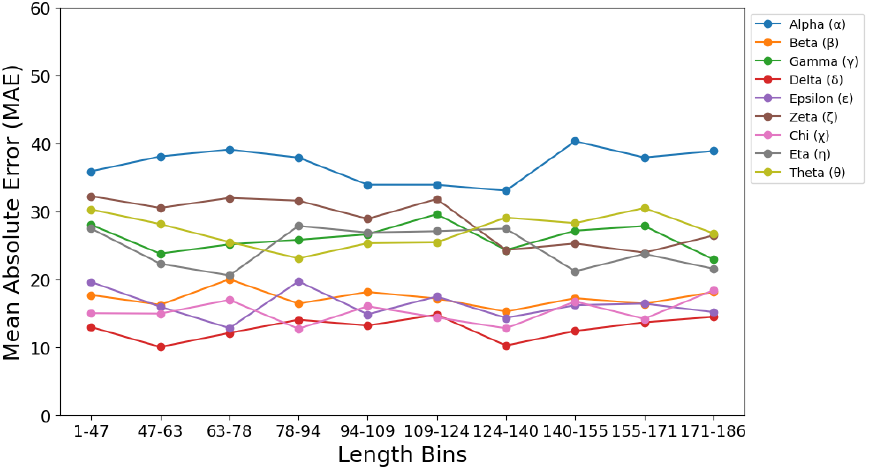
Variation of mean absolute error with respect to the length (number of nucleotides) of RNA sequences

### Performance for nucleotides in various pairing interactions

The performance of our model was evaluated on torsion angle prediction across different RNA interactions within the test datasets TS1, TS2, and TS3. Each dataset was analyzed based on the percentage of nucleotides involved in various structural interactions: unpaired bases, lone pairs, pseudoknots, multiplets, noncanonical pairs, and canonical nested base pairs. The predictive accuracy of our model was compared to the SPOT-RNA-1D as shown in Table 1.

Analysis revealed that nucleotides involved in tertiary interactions, such as lone pairs, pseudoknots, multiplets, and noncanonical pairs, consistently presented greater challenges in torsion angle prediction across all three test sets. This was evident from the consistently higher mean absolute errors (MAEs) for these interactions compared to canonical nested base pairs, which had the lowest MAEs, indicating their relative predictability due to less structural complexity. In TS1, the unpaired bases showed the most significant challenge in prediction due to more variability and flexibility in structure, with MAEs of 59.55 for *α*, considerably higher than those for canonical nested pairs, which had an MAE of 32.02 for the same angle. This pattern was consistent across all angles and other test datasets. TS2 and TS3 followed a similar trend, with slight variations in the percentages of nucleotides involved in each interaction type. For instance, TS2 had a slightly higher percentage of pseudoknots at 6.63%, compared to 5.8% in TS1, which contributed to the slightly varied prediction challenges observed. TS3, characterized by different experimental derivation techniques (NMR vs. X-ray crystallography for TS1 and TS2), still maintained the general trend where more complex tertiary interactions resulted in higher prediction errors.

Comparative analysis between our model and SPOT-RNA-1D [8] demonstrated that despite the inherent challenges posed by complex interactions, our model achieved consistently lower MAEs across most interactions and torsion angles, indicating an overall improvement in prediction accuracy. Our findings suggest that the complexity of nucleotide interactions significantly influences the difficulty of predicting torsion angles in RNA structures. More specifically, interactions that introduce greater structural complexity and variability tend to result in higher prediction errors. The comprehensive evaluation across multiple test sets confirms the robustness of our model, particularly in its superior performance over the baseline in handling the intricate and variable interactions within RNA structures.

### Performance on RNA-TorsionBERT dataset

Table 3 presents the performance of our method, DeepRNA-Twist, in predicting RNA torsion angles compared to various state-of-the-art approaches including RNA-TorsionBERT [9], SPOT-RNA-1D [8], and methods from the recent State-of-the-RNArt [20] benchmark on RNA-TorsionBERT dataset. DeepRNA-Twist demonstrates superior accuracy with the lowest mean MAE of 19.78° across all torsion angles. This represents a significant improvement over RNA-TorsionBERT [9], which records a mean MAE of 22.6°, and a more pronounced advantage over SPOT-RNA-1D with a mean MAE of 30.0°. Specifically, DeepRNA-Twist achieves the best performance in predicting *δ* with an MAE of 12.1°, surpassing RNA-TorsionBERT’s MAE of 13.6° and greatly outperforming SPOT-RNA-1D’s 28.3°. The largest improvement over RNA-TorsionBERT [9] is observed in the *γ* angle, where DeepRNA-Twist reduces the MAE by nearly 4°. Moreover, across other angles such as *χ, ϵ*, and *β*, DeepRNA-Twist consistently maintains lower MAEs compared to all benchmarked methods. We also notice the same trend in the relationship between prediction difficulty and the distribution of angles(Figure 4). Angles associated with wider distribution, such as *α, γ*, and *ζ*, are more difficult to predict compared to angles with narrower distribution such as *δ* and *ϵ*. In a nutshell, DeepRNA-Twist consistently shows statistically significant improvements in MAE across all torsion angles in this dataset over all other methods (Table 4).

## Ablation Study

To evaluate the contributions of different components within the DeepRNA-Twist architecture, we conducted an ablation study in two parts: (1) a comparison between RiNALMo embeddings and one-hot encoded features, and (2) an assessment of individual architectural components by systematically removing them. The results are summarized in Tables 5 and 6, which include results for both RNA-TorsionBERT and SPOT-RNA-1D datasets. We first compared the performance of DeepRNA-Twist model using RiNALMo embeddings against a simpler feature encoding approach with one-hot encoded nucleotides. As shown in Table 5, the model using RiNALMo embeddings consistently outperforms the one-hot encoded model on both RNA-TorsionBERT and SPOT-RNA-1D datasets. In the second part of our ablation study, we evaluated the contributions of key components within DeepRNA-Twist architecture by systematically removing them. Each component’s removal allowed us to assess its impact on the overall model performance, as reflected by the mean MAE of all torsion angles. Table 6 presents the results of the component removal experiments. The removal of the 2A3IDC modules led to the highest increase in mean MAE across both datasets, emphasizing their critical role in capturing both local and global dependencies within the sequence. Removing the Transformer Encoder Layer and Multi-head Attention also resulted in performance drops, highlighting their importance in the model’s architecture. The ablation study confirms that the 2A3IDC modules and language model embeddings are vital, validating our design choices in DeepRNA-Twist.

**Table 5.** Feature Encoding Ablation: Mean MAE of 9 Torsion Angles.

| (a) RNA-TorsionBERT Dataset |  |
| --- | --- |
| Feature Encoding | Mean MAE ($^{\circ}$) |
| **RiNALMo** | **19.78** |
| One-Hot Encoding | 23.45 |

| (b) SPOT-RNA-1D Dataset |  |
| --- | --- |
| Feature Encoding | Mean MAE ($^{\circ}$) |
| **RiNALMo** | **23.58** |
| One-Hot Encoding | 27.31 |

**Table 6.** Component Ablation: Mean MAE of 9 Torsion Angles.

| (a) RNA-TorsionBERT Dataset |  |
| --- | --- |
| Model Variant | Mean MAE ($^{\circ}$) |
| **Full Model** | **19.78** |
| Without Transformer Encoder | 22.90 |
| Without 2A3IDC Modules | 24.31 |
| Without Multi-head Attention | 21.86 |

| (b) SPOT-RNA-1D Dataset |  |
| --- | --- |
| Model Variant | Mean MAE ($^{\circ}$) |
| **Full Model** | **23.58** |
| Without Transformer Encoder | 26.34 |
| Without 2A3IDC Modules | 28.72 |
| Without Multi-head Attention | 25.91 |

## Case Study

In this case study, we utilized DeepRNA-Twist to predict the torsion angles of two RNA structures with PDB ID 7PTK and 4R4V, and compared to the native structures obtained from the PDB. The predicted torsion angles by DeepRNA-Twist were used to construct the predicted RNA structures, which were then superimposed on the original RNA structures (see Figure 6). In the superimposed visualizations, the native structures are displayed in cyan, while the predicted structures are shown in magenta, allowing for a direct comparison of the model’s predictions against the experimental data. We used PyMOL [30] for the visualization. The Root Mean Square Deviation (RMSD) values were calculated to quantify the accuracy of the predictions. For 7PTK, the RMSD between the predicted and native structures was 6.59 Å, and for 4R4V, the RMSD was 3.31 Å. These results demonstrate the model’s capability to closely align predicted RNA structures with their experimentally determined counterparts, highlighting the effectiveness of DeepRNA-Twist in RNA torsion angle prediction.

**Fig 6.**
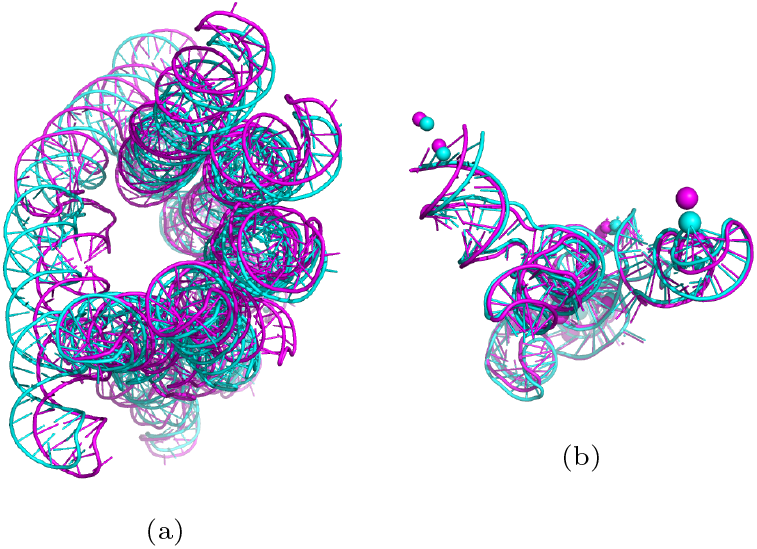
Visualization of the superposition of RNA structures: (a) 7PTK and (b) 4R4V using the torsion angles estimated by DeepRNA-Twist (magenta color), compared with the structures using the native angles obtained from PDB (cyan color).

## Discussion

In this study, we introduced DeepRNA-Twist, a novel deep learning approach for predicting RNA torsion and pseudo-torsion angles solely from sequence data. DeepRNA-Twist integrates several key modules, each contributing unique strengths. The transformer encoder captures complex dependencies and contextual relationships, essential for RNA torsion angle prediction. The 2A3IDC modules extract multi-scale features and recognize long-range dependencies while refining focus on critical structural variations through multi-head attention mechanisms. Attention modules throughout the architecture dynamically weigh contributions, ensuring significant features are prioritized. Our evaluations on benchmark datasets such as RNA-Puzzles [12], CASP-RNA [13], and SPOT-RNA-1D [8] demonstrate that DeepRNA-Twist significantly outperforms existing methods, achieving state-of-the-art accuracy. On the SPOT-RNA-1D dataset, the model showed 10% to 15% MAE improvements, depicting its robustness and accuracy. Angles with wider distribution like *α, ζ*, and *θ* showed notable improvements, whereas angles like *δ* and *ϵ*, which have narrower distribution, were easier to predict. Despite complexities in predicting angles for nucleotides involved in tertiary interactions, DeepRNA-Twist consistently achieved lower MAEs compared to other methods. Further analysis on RNA-Puzzles and CASP-RNA datasets highlights that DeepRNA-Twist outperforms deep learning based state-of-the-art methods, including RNA-TorsionBERT [9], SPOT-RNA-1D [8], and AlphaFold 3 [22], with an overall MAE reduction of more than 10° and a 35% improvement in accuracy compared to template-based and ab initio methods. These results highlight the model’s superior performance and generalizability. Despite these advancements, inherent limitations remain. Reducing MAE below a certain threshold is challenging due to RNA structural complexity and variability. Angles like *α* and *γ* with wider distribution are particularly difficult to predict accurately, necessitating continued research, the integration of additional biophysical constraints and more comprehensive datasets. In our future work, we aim to address these issues to further improve prediction accuracy.

